# Dichotomous Activity and Function of Neurons with Low- and High-Frequency Discharge in the External Globus Pallidus of Non-Human Primates

**DOI:** 10.1101/2021.12.19.473353

**Authors:** Shiran Katabi, Avital Adler, Marc Deffains, Hagai Bergman

**Affiliations:** Department of Medical Neuroscience, Institute of Medical Research Israel-Canada (IMRIC), The Hebrew University-Hadassah Medical School, 91120 Jerusalem, Israel; The Edmond and Lily Safra Center for Brain Sciences, The Hebrew University, 91904 Jerusalem, Israel; University of Bordeaux, UMR 5293, IMN, 33000 Bordeaux, France; CNRS, UMR 5293, IMN, 33000 Bordeaux, France; Department of Neurosurgery, Hadassah Medical Center, 91120 Jerusalem, Israel

## Abstract

To date, there is a consensus that there are at least two neuronal populations in the non-human primate (NHP) external globus pallidus (GPe): the low- and high-frequency discharge (LFD and HFD) neurons. Nevertheless, almost all NHP physiological studies have neglected the functional importance of LFD neurons. This study examined the discharge features of these two GPe neuronal subpopulations recorded in four NHPs engaged in a classical conditioning task with cues predicting reward, neutral and aversive outcomes. The results show that LFD neurons tended to burst, encoded the salience of behavioral cues, and exhibited correlated spiking activity. By contrast, the HFD neurons tended to pause, encoded cue valence, and exhibited uncorrelated spiking activity. Overall, these findings point to the dichotomic organization of the NHP GPe which is likely to be critical to the implementation of normal basal ganglia functions and computations.

## Introduction

The globus pallidus (GPe) is a central nucleus in the basal ganglia (BG) circuitry and a key player in BG computational physiology and pathophysiology ^1–3^. At least two different neuronal populations are known to exist within the GPe. Earlier in vivo electrophysiological recordings in rodents ^4^, non-human primates (NPHs) ^5^ and humans ^6^ indicated that GPe neurons can be defined as a function of their specific discharge rates and patterns into: low-frequency discharges with occasional brief high frequency bursts (LFD neurons) and high-frequency discharges occasionally interrupted by pauses (HFD neurons). However, most computational models of the BG treat the GPe as a single GABAergic cell-type nucleus ^7–9^.

Other studies using a juxtacellular recording-labeling technique in dopamine-depleted ^10^ and dopamine-intact ^11^ anesthetized rats have identified two categories of GABAergic GPe neurons. These two categories of GPe neurons, termed “arkypallidal” and “prototypic” neurons, exhibit distinct neurochemical, structural and electrophysiological features. The arkypallidal neurons provide exclusive feedback projections to the striatum and parallel the well-known feedback projections from GPe to subthalamic nucleus (STN). They target both striatal projection neurons and interneurons. The prototypic neurons project to the STN and BG output structures. Recent studies have also shown that the arkypallidal and prototypic neurons are differentially impacted by striatal and STN inputs ^12,13^, thus suggesting separated functional pathways in the rodent GPe. In line with this view, the role of GPe neuronal subpopulations in regulating movement was reported to be dissociable in behaving rodents ^14,15^. Specifically, the arkypallidal neurons exhibited stronger and faster responses to a Stop signal than the prototypic neurons, coherent with the suppression of the development of the Go-related striatal activity ^14^. Given the differences in rate and pattern of these two neuronal subpopulations ^10,11,14^, it is likely that the NPH GPe LFD and HFD neurons represent the rodent GPe arkypallidal and prototypic neurons, respectively. Anatomical studies confirm significant (15% of labeled GPe neurons) back projections from the GPe to the striatum also in NHPs ^16^. Given the unique value of the NHP model in translational research, it is of utmost importance to explore the dichotomous organization in the NHP GPe to determine its role in BG information processing. To tackle this challenge, we examined the discharge (rate and pattern) features, behavioral responses, as well as the signal and noise correlations of LFD and HFD neurons recorded in the GPe ^17^ of four NHPs engaged in a classical conditioning task with cues predicting reward, neutral and aversive outcomes (Fig. 1a).

**Figure 1.**
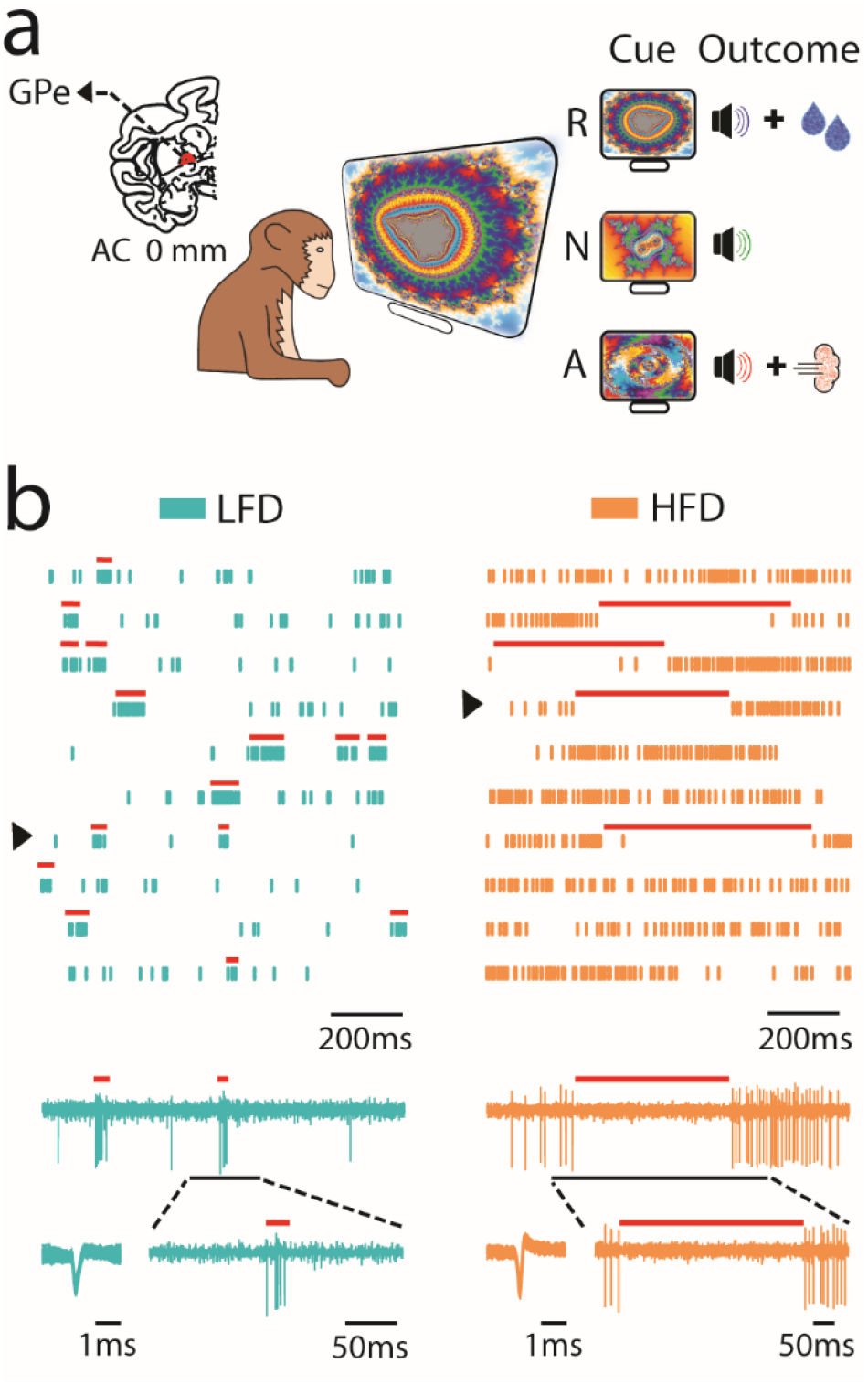
Behavioral task and neuronal recordings. **a**, Classical conditioning paradigm and recording site. Different visual cues were introduced to the monkey for a 2-s period which predicted the probability or the intensity of food (R, reward), neutral (N), or air-puff (A, aversive) outcomes. All outcome deliveries were signaled by one of three sounds and were followed by a variable intertrial interval (5-6 s). Top left, Representative coronal section 0 mm from the anterior commissure (AC 0), adapted from Martin and Bowden ^17^, shows the external globus pallidus (GPe) recording site. **b**, Examples of low-frequency discharge (LFD) and high-frequency discharge (HFD) neuronal activities. Raster plots with marked LFD bursts (left) and HFD pauses (right) as detected by optimal burst/pause detection automatic methods (see Methods). Rasters show 10 consecutive 1-s segments (from bottom to top and from left to right). An example of the multi-unit activity datasets filtered between 250 and 6000 Hz of a 1-s segment (marked by an arrow) appears beneath each raster plot. Beneath the 1-s segment, one of the bursts (left) and one of the pauses (right) are presented on a smaller time scale along with their examples of spike waveforms. The spike waveform plot is composed of 100 superimposed waveforms of randomly selected spikes from the whole recording period of the neuron.

## Results

### LFD and HFD neurons exhibit different discharge features

GPe neurons were defined online as LFD and HFD neurons (n = 98 and 632 neurons, respectively) according to their discharge rate and pattern. Exemplary raster plots of the spike trains and raw analog data of the LFD and HFD neurons are depicted in Fig. 1b. LFD neurons were previously reported to fire between 1 and 30 spk/s (mean = 10 spk/s), whereas HFD neurons fire between 10 and 100 spk/s (mean = 55 spk/s) ^5^. Similarly, in our database, the firing rate of the LFD neurons was significantly lower than the firing rate of the HFD neurons (mean ± SEM = 13.6 ± 1.08 and 71.16 ± 1.1 spk/s, respectively, Mann-Whitney U-test, p < 0.001; Fig. 2a). In addition, HFD neurons had a significant narrower waveform length (calculated as the duration from the first negative peak to the next positive peak of the extracellularly recorded action potential) than LFD neurons (mean ± SEM = 0.31 ± 0.003 and 0.83 ± 0.19 ms, respectively, Mann-Whitney U-test, p < 0.001; Supplementary Fig. 1a).

**Figure 2.**
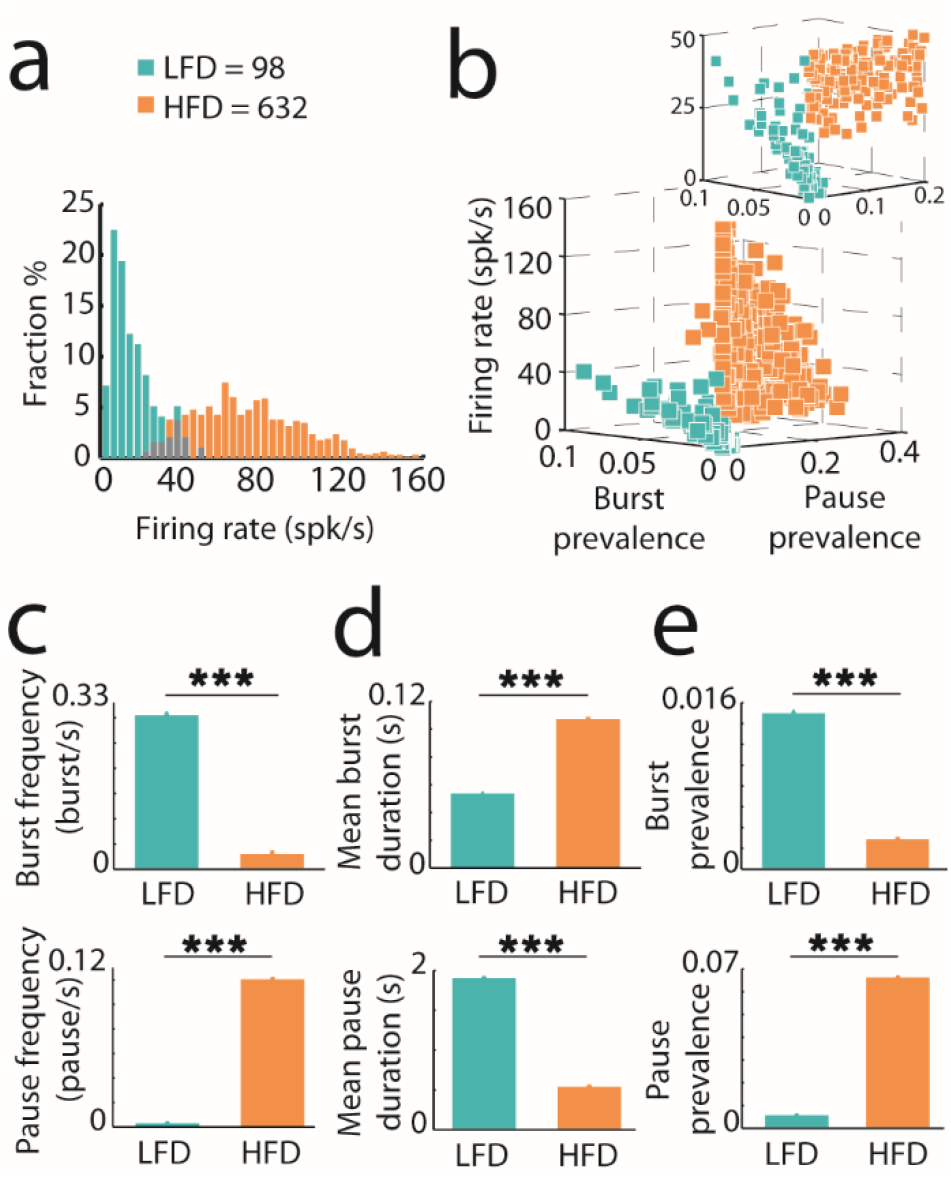
Dichotomous activity of identified GPe neurons. **a**, Histogram of the distribution of the firing rate of GPe neurons (bin width = 4 spk/s). **b**, Three-dimensional scatter plot of GPe neurons based on their firing rates and burst/pause prevalence values (LFD = 98, HFD = 632). Inset only depicts neurons with firing rate ≤ 50 spk/s (LFD = 98, HFD = 156). Each square represents a single neuron colored according to type. **c-e**, Means (± SEMs) of the frequency, duration and prevalence of the bursts and pauses in the spiking activity of the LFD and HFD neurons. ^***^ indicates significant (p < 0.001) differences between LFD and HFD neurons.

The mean interspike interval (ISI) of the LFD neurons (mean ± SEM = 75.09 ± 0.26 ms; Supplementary Fig. 1b, top) was ∼5.5-fold higher than the mean ISI of the HFD neurons (mean ± SEM = 13.51 ± 0.003 ms; Supplementary Fig. 1b, bottom), possibly reflecting the bursting pattern of the LFD neurons. In line with previous publications ^5^ and the bursting pattern of the LFD neurons (see below), the modal ISI was shorter for the LFD neurons than the HFD neurons (Supplementary Fig. 1b). The mean coefficient of variance of the ISIs (CV ISIs, a measure of the variability of the firing pattern) of the LFD neurons (mean ± SEM = 2.97 ± 0.17; Supplementary Fig. 1c, top) was significantly higher than the mean CV ISIs of the HFD neurons (mean ± SEM = 1.75 ± 0.02; Supplementary Fig. 1c, bottom; Mann-Whitney U-test, p < 0.001), thus pointing to the more irregular and bursting firing pattern of the LFD neurons^18^.

Auto-correlograms of the neurons were also constructed. The auto-correlation function depicts the distribution of times between any two spikes (not only successive as in the ISI histogram) in a spike train and therefore is particularly useful for detecting specific discharge patterns ^19^. The averaged auto-correlogram of the LFD neurons, like their ISI histogram, had an earlier and sharper peak (mean ± SEM = 6.98 ± 0.11 ms; Supplementary Fig. 1d, top) than the peak of the averaged auto-correlogram of HFD neurons (mean ± SEM = 35.41 ± 0.55 ms; Supplementary Fig. 1d, bottom). This early sharp peak in the auto-correlogram is a characteristic of the bursting firing pattern of the neurons ^19^. Short-term peaks in the auto-correlation functions can also be the result of the refractory period of cells with high firing rate ^20^. However, this was probably not the case for the LFD neurons here due to their relatively low firing rates (< 50 spk/s) and the combination of additional measures that reflect the bursting firing of these neurons (See below).

### LFD tendency to burst vs. HFD tendency to pause

For burst/pause detection, we applied Poisson surprise algorithms (See Methods and Fig. 1b). Since the firing rate distributions of the online identified LFD and HFD neurons revealed a certain degree of overlap between ∼20 and ∼50 spk/s (Fig. 2a), we generated a three-dimensional scatter plot of all GPe neurons based on their firing rates and burst/pause prevalence values (Fig. 2b). We then subjected the neurons with firing rates ≤ 50 spk/s to further offline binary classification. We performed 3D k-means cluster analysis of all spike trains with k = 2 clusters, using the discharge rates, burst prevalence values (i.e., the probability of bursts of spikes in a detected spike train) and pause prevalence values (i.e., the probability of spike-free long segments in a detected spike train). The binary classification of each cluster was performed and a confusion matrix (also known as an error matrix) was built to determine to what extent the online identified LFD and HFD neurons belonged to each cluster. The true negative rate (specificity) and true positive rate (sensitivity) were high (92.31% and 84.69%, respectively), indicating that misclassification of the online identified LFD and HFD neurons within each cluster was relatively low, and therefore validating the use of the online identified neurons for further analyses. Equally important, these results revealed that LFD and HFD neurons with similar firing rates could be efficiently discriminated based on their burst and pause features.

We further averaged the frequency, mean duration and prevalence (frequency * mean duration) of bursts/pauses for all LFD and HFD neurons (Fig. 2c-e). We found that the mean burst prevalence was significantly higher in the LFD neurons than the HFD neurons, and vice-versa for pause prevalence (Mann-Whitney U-test, p < 0.001). Furthermore, the fraction of LFD neurons exhibiting bursts was systematically higher than the fraction of HFD neurons (Supplementary Fig. 2a). Contrariwise, the fraction of HFD exhibiting pauses was higher than the fraction of LFD neurons (Supplementary Fig. 2b). Altogether, these results indicate that LFD and HFD neurons represent two neuronal populations with distinct discharge rates and patterns.

Finally, in line with previous publications that LFD bursts usually consist of 5-20 spikes ^5^, LFD neurons had bursts with 10.76 ± 0.04 spikes (Supplementary Fig. 3a, top) and a mean duration of 49.77 ± 0.29 ms (Supplementary Fig. 3b, top). The averaged intraburst ISI pattern of all LFD and HFD neurons for different burst lengths revealed that the duration of intraburst ISIs of the bursts of LFD neurons decreased during the first half of the burst and increased towards the end of the burst (Supplementary Fig. 3c, top). This intraburst ISI pattern of LFD neurons has been described as the accelerating-decelerating (parabolic) burst ^21,22^. In contrast, this type of pattern was not found for the HFD neurons (Supplementary Fig. 3c, bottom), which had longer bursts with 18.28 ± 0.07 spikes (Supplementary Fig. 3a, bottom) and a mean duration of 97.42 ± 0.56 ms (Supplementary Fig. 3b, bottom).

### LFD and HFD neurons exhibit dichotomic task-related activity

Examination of the GPe population responses to cues with different motivational/emotional values predicting liquid rewards (reward cues), air-puffs to the eyes (aversive cues) or neither (neutral cues), revealed that the LFD and HFD neurons exhibited diverse response profiles. LFD neurons displayed homogenous transient population responses followed by low persistent activity to all cues (Fig. 3a, top). By contrast, HFD neurons displayed persistent and opposite population responses to reward and neutral/aversive cues (Fig. 3b, top). Reward, neutral and aversive cue-related activities (i.e., mean values of the activity during the 2 s duration of reward, neutral and aversive cues, respectively) differed significantly from baseline activity (i.e., mean values of the activity during baseline) for LFD neurons (Wilcoxon signed rank tests, p < 0.001, for each cue; Fig. 3a, top). However, there was no significant difference between their different cue-related activities (Friedman test, p = 0.22; Fig. 3a, bottom). By contrast, only reward cue-related activity differed significantly from baseline activity for HFD neurons (Wilcoxon signed rank tests, p < 0.001, p = 0.18 and p = 0.14, for reward, neutral and aversive cue-related activities, respectively; Fig. 3b, top). Accordingly, we found a significant effect of cue value on cue-related activity for HFD neurons (Friedman test, p < 0.001), such that reward cue-related activity differed significantly from both neutral and aversive cue-related activities (post hoc comparisons, Bonferroni corrected, p < 0.001, for both reward and neutral or reward and aversive cues; Fig. 3b, bottom). To better emphasize the opposite polarity of the early population response of the HFD neurons to reward and neutral or aversive cues (Fig. 3b, top), we also ran these analyses using a 0.5 s cue duration (instead of 2 s) for the calculation of the cue-related activities (Supplementary Fig. 4a-b) and yielded similar results.

**Figure 3.**
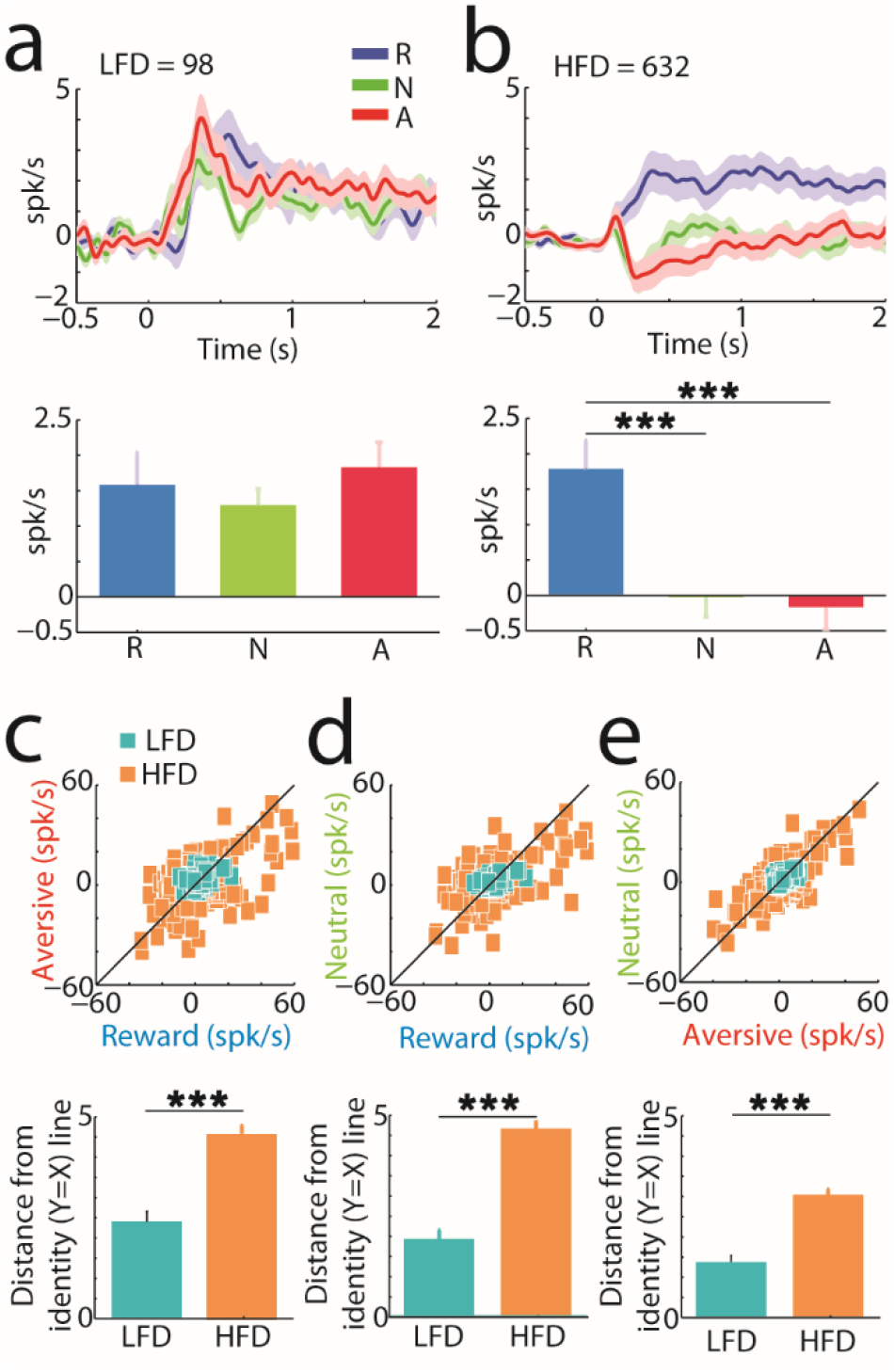
Dichotomous function of LFD and HFD neurons in motivational/emotional cue processing: salience vs. valence. **a-b**, Top, LFD and HFD population responses to behavioral cues presented at time 0. The responses of a single neuron to the cues were characterized by their PSTHs (calculated in 1-ms bins, smoothed with a Gaussian window with a SD of 20 ms, and normalized by subtracting their baseline firing rate, i.e., mean firing rate in the last 0.5 s of the intertrial interval). Population responses to the cues were defined as the averaged responses of all neurons. The shaded areas mark the SEMs. R, responses to reward cues; N, responses to neutral cues; A, responses to aversive cues. Bottom, mean LFD and HFD reward, neutral and aversive cue-related activities. Cue-related activity is defined as the mean value of the activity for the 2 s cue duration. Error bars represent SEMs; ^***^ indicates significant (p < 0.001) differences between cue-related activities. **c-e**, Top, comparison between reward/aversive, reward/neutral and aversive/neutral cue-related activities. Each square refers to a single neuron. The black line is the identity (Y = X) line. Bottom, mean distance values of the squares from the identity line. Error bars represent SEMs; ^***^ indicates significant (p < 0.001) differences between LFD and HFD neurons.

Next, we compared the reward/aversive, reward/neutral and aversive/neutral cue-related activities of the LFD and HFD neurons (Fig. 3c-e and Supplementary Fig. 4c-e). In line with the dichotomic response profile of the LFD and HFD populations to incentive cues, the mean distance values of the cue-related activities from the identity line of the LFD neurons were significantly lower than the HFD neurons for each cue combination (Mann-Whitney U-tests, p < 0.001).

To quantify the differences between single-cell responses to reward and aversive cues vs. their response to the neutral cue, we calculated the response index for reward and aversive cues (i.e., the absolute difference between reward or aversive cue-related activity and neutral cue-related activity). As already reported ^23,24^, most of the LFD and HFD neurons exhibited a larger response index for the reward cues than for the aversive cues (64% and 68% of the LFD and HFD neurons, respectively; Supplementary Fig. 5a, top). However, the mean distance values from the identity line of the LFD neurons were significantly lower than the HFD neurons (Mann-Whitney U-test, p < 0.001; Supplementary Fig. 5a, bottom). The same analysis using a 0.5 s cue duration for the calculation of the cue-related activities produced similar results (Supplementary Fig. 5b). These results therefore indicate that the LFD neurons exhibit more similar responses to the behavioral cues than the HFD neurons.

Signal correlation measures the similarity of pairwise responses to behavioral events ^25–27^. In line with the single-cell results, we found that the SC distribution for LFD/LFD neuron pairs (Fig. 4a) tended toward positive values, indicating a tendency toward similar cue-related responses. By contrast, the SC distributions for HFD/HFD (Fig. 4b) and LFD/HFD (Fig. 4c) neuron pairs were more symmetrical, with an average SC closer to zero, indicating uncorrelated HFD neuronal responses to the cue.

**Figure 4.**
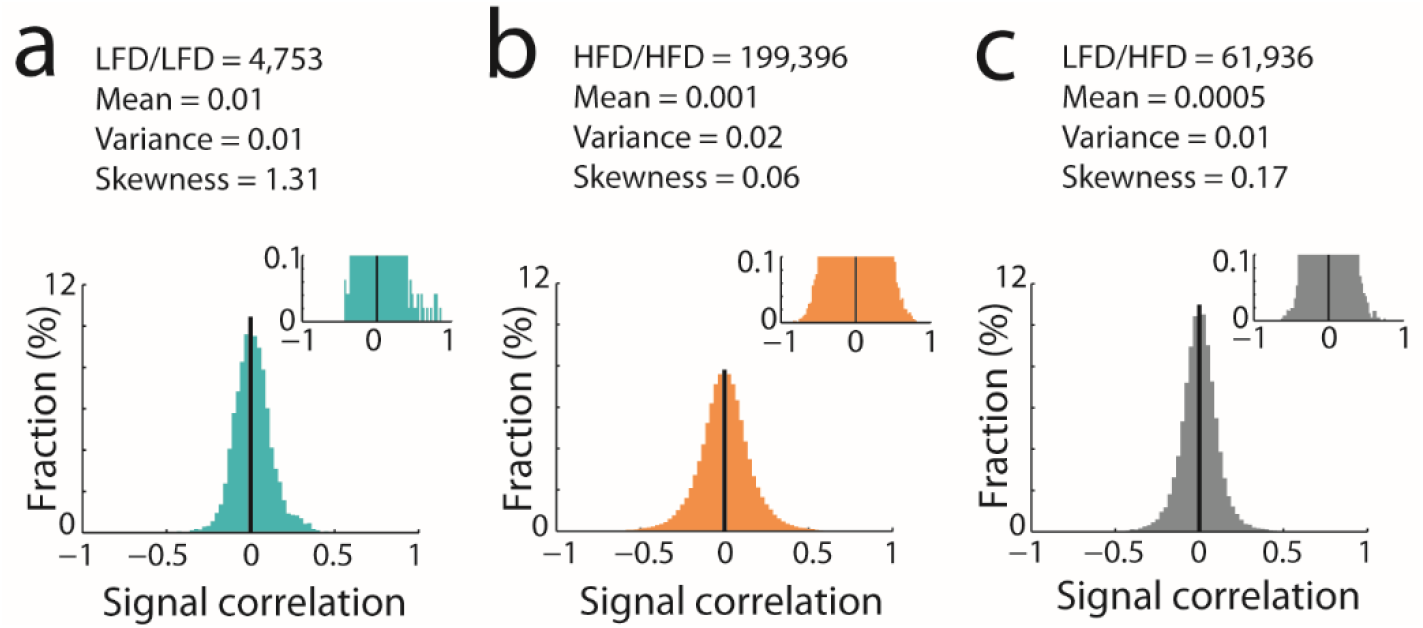
Higher similarity of the LFD task-related responses to the behavioral cues. **a-c**, Distribution of LFD/LFD, HFD/HFD and LFD/HFD pairwise signal correlations. In the insets, the y-scale was truncated to better depict the distribution tails. Mean, variance and skewness values are the features of the signal correlation (SC) distributions.

### LFD neurons exhibit spontaneous correlated spiking activity

The mean spike-to-spike (noise) cross-correlogram (CCH) of LFD/LFD neuron pairs exhibited a robust peak around zero (the mean value of the CCHs at time = 0, T_0 =_ 9.42 spk/s, Fig. 5a) indicative of a significant level of correlated activity between simultaneously recorded LFD neurons. This peak did not appear in the mean CCHs of HFD/HFD (T_0_ = 0.24 spk/s, Fig. 5b) and LFD/HFD (T_0_ = 0.22 spk/s, Fig. 5c) neuron pairs. Here, we corrected the CCHs by subtracting the shuffle predictor to remove any stimulus-related relationship. Therefore, we ruled out the possibility that correlated activity between simultaneously recorded LFD neurons was due to the homogeneity of the responses of the LFD neurons to the cues. Importantly, these results were maintained after recalculation (see Methods) of the T_0_ values of the mean corrected CCHs after random selection of a smaller number (20 pairs) of HFD/HFD and LFD/HFD neuron pairs (Fig. 5b,c, insets) similar to the small number of LFD/LFD neuron pairs (Fig. 5a). Therefore, the correlated activity between simultaneously recorded LFD neurons was not due to the confounding effects of the small number of LFD/LFD neuron pairs.

**Figure 5.**
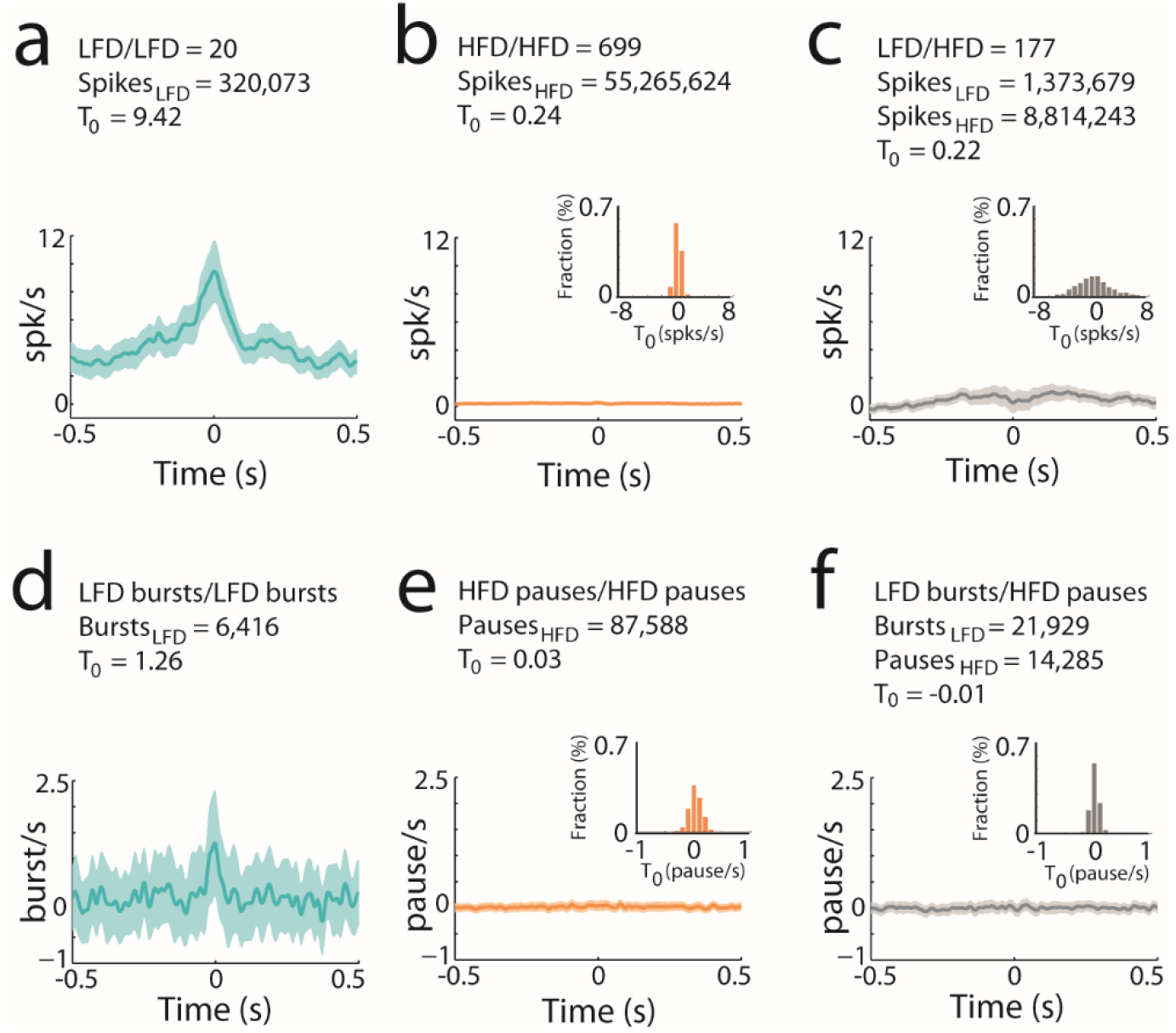
LFD activity is driven by a common input. **a-c**, Average spike-to-spike cross-correlograms for LFD/LFD, HFD/HFD and LFD/HFD neuron pairs. Each cross-correlogram was corrected by subtracting its shuffle predictor in bins of 1 ms and then averaged over all pairs. The shaded areas mark SEMs; the T_0_ value is the firing rate at time = 0; insets are the distributions of the T_0_ values of the mean corrected cross-correlograms of 1000 randomly selected 20 HFD/HFD and LFD/HFD neuron pairs (to match the number of LFD/LFD neuron pairs). **d-f**, Average burst-to-burst, pause-to-pause and burst-to-pause cross-correlograms of LFD/LFD, HFD/HFD and LFD/HFD neuron pairs, respectively. Each cross-correlogram was corrected by subtracting its shuffled predictor in bins of 1 ms and then averaged over all LFD bursts or HFD pauses. The shaded areas mark SEMs; the T_0_ value is the mean frequency of LFD bursts or HFD pauses at time = 0 (i.e., LFD burst or HFD pause initiation); Insets as in **b-c**.

Given these results, we examined whether the LFD neuron pairs’ bursts tended to coincide. We computed the mean burst-to-burst CCH of LFD/LFD neuron pairs and compared it to the mean pause-to-pause and burst-to-pause CCHs of HFD/HFD and LFD/HFD neuron pairs, respectively. This identified a coincidence between bursts detected in simultaneously recorded LFD/LFD neuron pairs (T_0_ = 1.26 burst/s; Fig. 5d). Conversely, the mean pause-to-pause and burst-to-pause CCHs of HFD/HFD and LFD/HFD neuron pairs remained flat (T_0_ = 0.03 and - 0.01 pause/s, respectively; Fig. 5e,f), even when recalculating the T_0_ values after random selection of only 20 neuron pairs of HFD/HFD and LFD/HFD (Fig. 5e,f, insets). Moreover, an increase in the LFD firing rate concurred with bursts detected in simultaneously recorded LFD neurons (T_0_ = 14.03 spk/s; Supplementary Fig, 6a), but not with pauses detected in simultaneously recorded HFD neurons (T_0_ = -0.12 spk/s; Supplementary Fig. 6d). Finally, HFD neurons presented a tendency toward a slight decrease in their firing rate in response to simultaneous HFD pauses (T_0_ = -1.03 spk/s; Supplementary Fig. 6b) or LFD bursts (T_0_ = -1.01 spk/s; Supplementary Fig, 6c). Altogether, these results provide strong evidence for LFD correlated spontaneous spiking activity in the GPe of healthy animals.

### LFD neuron pairs display a sustained noise correlation during behavioral cue periods

To further characterize the dynamics of LFD correlated activity during the behavioral cues, we compared the joint peri-stimulus time histograms (JPSTHs) ^25,26,28^ of the LFD/LFD, HFD/HFD and LFD/HFD neuron pairs (Fig. 6). We found that the LFD/LFD neuron pairs showed a persistent increase in noise correlation throughout the cue duration (Fig. 6a, top). The modulations (after baseline normalization) along the JPSTH main diagonal of the LFD/LFD neuron pairs differed between the cue duration (i.e., mean values of the main diagonals during the 2 s cue period) and the baseline period (i.e., mean values of the main diagonal during the last 0.5 s of the intertrial interval, Wilcoxon signed rank test, p < 0.05; Fig. 6a, bottom). Modulations along the JPSTH main diagonal did not significantly differ between the cue duration and the baseline for the HFD/HFD and LFD/HFD neuron pairs (Wilcoxon signed rank tests, p = 0.54 and p = 0.63, respectively; Fig. 6b,c, bottom). Nevertheless, the modulations along the normalized JPSTH main diagonal of the LFD/LFD neuron pairs were not significantly different from those of the HFD/HFD and LFD/HFD neuron pairs (Kruskal-Wallis H-test, p = 0.35; Fig. 6, bottom). Finally, although the dynamics of the LFD/LFD correlated activity seemed to vary across the behavioral cues (Supplementary Fig. 7), these modulations along the main diagonals of the different types of neuron pairs were not significantly different between the reward, neutral and aversive cue periods (Friedman tests, p = 0.057, p = 0.87 and p = 0.8 for LFD/LFD, HFD/HFD and LFD/HFD neuron pairs, respectively). These results therefore provide compelling arguments for a sustained noise correlation, with no cue-related dynamics, of the LFD/LFD neurons throughout the behavioral task.

**Figure 6.**
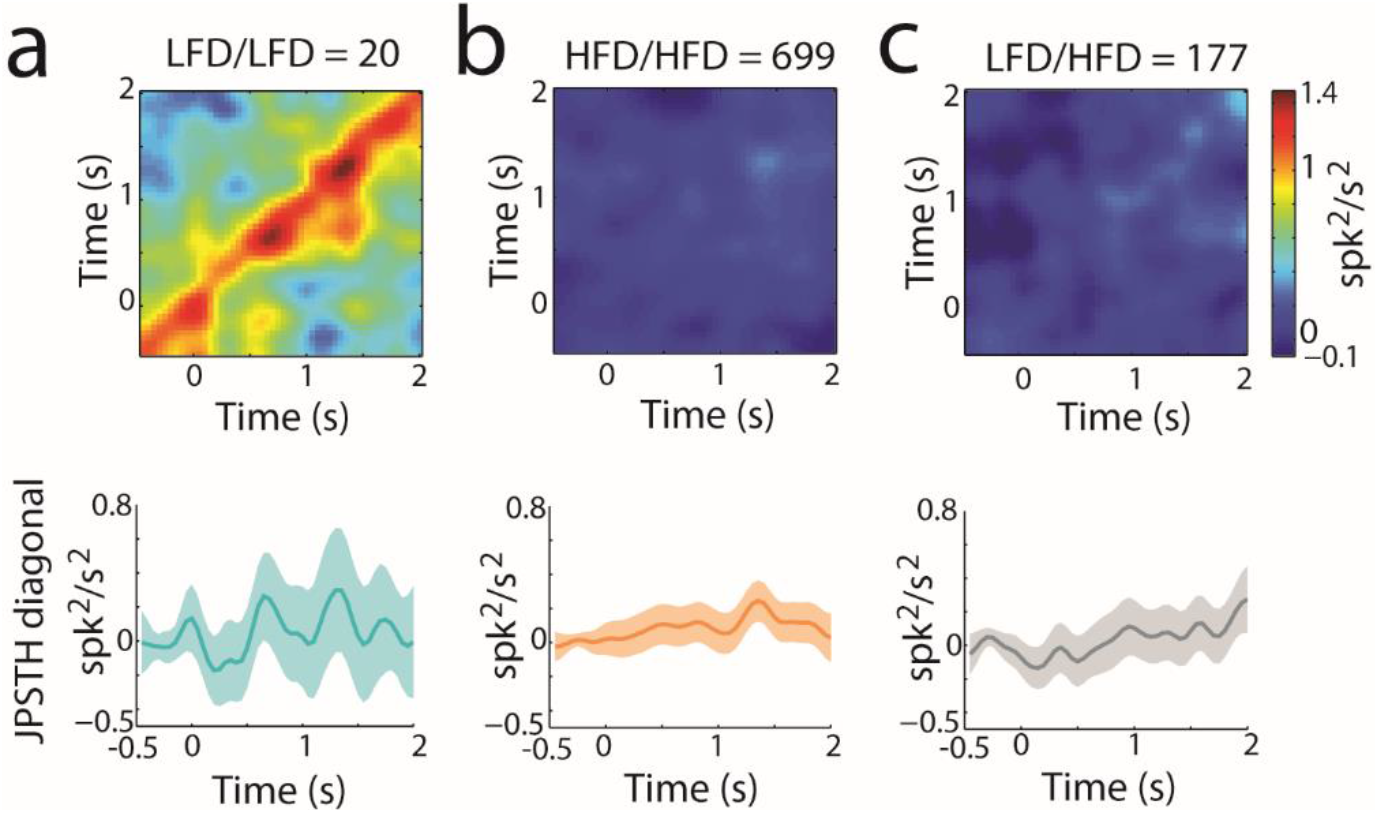
Sustained noise correlation of the LFD neurons during behavioral cue periods. **a-c**, Top, Population JPSTHs of LFD/LFD, HFD/HFD and LFD/HFD neuron pairs for behavioral cues presented at time 0. Each JPSTH was corrected by subtracting its JPSTH predictor in bins of 50 ms, smoothed with two-dimensional Gaussian window (SD = 50 ms) and then averaged over all pairs. The JPSTHs have the same color scaling (color bar on the right) to enable comparisons between the different types of neuronal pairs. Bottom, JPSTH main diagonals (± SEM, shaded envelop) for LFD/LFD, HFD/HFD and LFD/HFD neuron pairs. The main JPSTH diagonal of each pair was normalized by subtracting the mean value of the last 0.5 s of the intertrial interval, and then averaged across the whole population.

## Discussion

We examined the discharge (rate and pattern) features and the responsiveness to incentive cues of LFD and HFD neurons recorded in the NHP GPe (n = 98 and 632, respectively, from four NHPs). In line with early descriptions ^5^, we found that LFD neurons only accounted for 13% of all recorded GPe neurons. Most of the recorded GPe neurons (87%) consisted of HFD neurons. We characterized the spontaneous and evoked discharge patterns and synchronizations of the two GPe neuronal populations and identified a dichotomous organization of the NHP GPe nucleus. The LFD neurons tended to burst, encoded the salience of behavioral cues, and evidenced synchronized activity. By contrast, the HFD neurons tended to pause, encoded the valence of behavioral cues, and exhibited independent (uncorrelated) spiking activity.

### LFD neurons tend to burst whereas HFD neurons tend to pause

Bursting activity in the GPe has been reported in rodents ^4^, cats ^29^, NHPs ^5^ and humans ^6^. In contrast to the extensive research on the pause properties of the HFD neurons ^30–32^, very little is known about the burst properties of LFD neurons. Here, we used the Poisson surprise algorithms for burst/pause detection. Our results revealed a higher probability of bursts for the LFD neurons than for the HFD neurons and a higher probability of pauses for the HFD neurons than for the LFD neurons (Fig. 2c-e). We also found that the fraction of cells defined as bursters and pausers depended strongly on the parameters used for their identification (e.g., percentage of 1-min segments of spiking activity with at least n bursts; Supplementary Fig. 2). Nevertheless, regardless of the parameter used for the definition of burster and pauser cells, the fraction of the so-called LFD bursters was systematically higher than the fraction of the so-called HFD bursters (Supplementary Fig. 2a), and vice-versa for the fraction of the so-called LFD and HFD pausers (Supplementary Fig. 2b). Finally, it is noteworthy that using the same threshold value for the identification of the pausers cells as reported in Elias *et al*. ^30^ (i.e., at least 80% of 1-min segments of spiking activity with at least two pauses) generated similar results of ∼50% pausers out of all the HFD neurons (Supplementary Fig. 2b, right). Therefore, our findings revealed a clear dichotomy in the discharge rate (lower vs. higher) and pattern (bursts vs. pauses) of the activity of LFD and HFD neurons recorded in the same animals and under the same experimental conditions.

### Functional bipartite organization of pallidal neurons

The recent discovery in the rodent that actions-in-preparation might be specifically cancelled via the feedback GABAergic projections from the GPe to the striatum ^1,14,15^ strongly suggests a specific physiological action of the arkypallidal neurons compared to the prototypic neurons in movement control. Nevertheless, the role of the GPe is not limited to motor control. Rather, GPe activity can also be driven by cues predicting future reward probability ^33^ and fluctuates differently in response to appetitive and aversive events ^24^. A more recent study in monkeys performing a deterministic reversal task also showed that GPe activity predicted whether the animals would switch or maintain their previous choice ^34^. Finally, a subset of GPe neurons that innervate the parafascicular intra-laminar nucleus of the thalamus has been identified in rodents and shown to be involved in behavioral flexibility ^35^. Nonetheless, despite the clear evidence for the implication of the GPe in non-motor processes and especially in information processing of incentive sensory cues, the specific contribution of the two pallidal subpopulations (NHP LFD/HFD neurons and rodent arkypallidal/prototypic neurons) remains unexplored.

In line with earlier NHPs studies ^24,36^, we found that HFD neurons exhibited persistent and uncorrelated activity in response to behavioral events, which mainly consist of pronounced increases in activity in response to reward cues and relatively small decreases in activity following aversive and neutral cues (Figs. 3b and 4b, and Supplementary Fig. 4b). Our results therefore demonstrate that HFD neurons asymmetrically encode the positive and negative valences of incentive cues, and contribute to the large information capacity of the BG network ^37^. An earlier study also reported that the activity of low-frequency discharge BG neurons (including the striatal projection neurons and the pallidal LFD neurons) was more strongly modulated by expectation of the reward than by expectation of the aversive event ^23^. However, when compared to the persistent and heterogeneous responses of the HFD neurons (Figs. 3b and 4b, and Supplementary Fig. 4b), we found that LFD neurons exhibited more phasic and correlated responses and did not efficiently discriminate between reward and non-rewarding cues (Figs. 3a and 4a, and Supplementary Fig. 4a). Therefore, these transient and more homogeneous responses of the LFD neurons most likely correspond to the salience encoding concept.

Interestingly, similar neuronal heterogeneity has been recently observed in the ventral pallidum (VP) of NHPs ^38^. In fact, it has been suggested that “transient” VP neurons might be involved in the estimation of action outcomes (the “critic-like function” in machine learning terminology), whereas “persistent” VP neurons would contribute to the control of the action (the “actor-like function”). This could imply that GPe LFD and HFD neurons correspond to the “transient” and “persistent” VP neurons, respectively. In line with this supposition, LFD neurons exhibited homogenous responses (Fig. 4a) similar to the homogeneous responses of the BG neuromodulators/critics (i.e., midbrain dopaminergic neurons and striatal cholinergic interneurons) ^39,40^, but contrasted with the heterogeneous responses of neurons in the BG main axis/actor system that include the HFD neurons (Fig. 4b) ^25,36^.

### Correlated pallidal activity in healthy state

Understanding the correlational features of normal (healthy) GPe activity is crucial, especially since emergence of correlated activity in the GPe is considered to be one of the main hallmarks of Parkinson’s disease ^41^. In the current study, we did not observe correlated activity between the simultaneously recorded HFD/HFD neurons of healthy NHPs (Fig. 5b). However, as previously reported ^32^, a slight decrease in HFD firing rate (∼ 1 spk/s) was found when averaging the simultaneous firing activity of HFD neurons over all pauses (Supplementary Fig. 6b). This effect, which was too small to be identified in a pairwise cross-correlation analysis that was limited to specific neuron pairs, suggests that the tendency of the HFD neurons to pause and/or decrease their firing rate might be network-driven ^32^. In any case, the synchronization of the HFD neuron pairs was weak, if not actually nonexistent, compared to the synchronization of the LFD neuron pairs. We found a robust peak around zero in the mean spike-to-spike CCH of the LFD/LFD (Fig. 5a). Moreover, the LFD bursts tended to occur together (Fig. 5d) and to coincide with a significant increase in firing rate of other LFD neurons (i.e., while a single LFD neuron is bursting, others are also activated, Supplementary Fig. 6a).

Remarkably, this LFD/LFD correlated activity was sustained during cue-related processing, even when comparing the different cue periods (reward, neutral and aversive), thus revealing a constant common input to the LFD neurons during over-trained behavior (Fig. 6a and Supplementary Fig. 7a). Earlier studies using CCHs/JPSTHs ^25,26^ have reported dynamics in the correlated spiking activity across the BG network (e.g., midbrain dopaminergic neurons and putamen medium spiny neurons), signifying a different encoding of the cue value in terms of neuronal synchronization. However, our results excluded such valence encoding of incentive cues by the LFD neurons. Taken together, these results therefore reveal distinct functional connectivity and response patterns for the two GPe subpopulations that might underlie their respective functional role in BG information processing.

Consistently, recent studies in rodents ^12,13^ have shown that although the striatum and STN massively innervate both arkypallidal and prototypic neurons, prototypic neurons only receive very weak inputs from the direct pathway striatal (D1) projection neurons ^13^. In addition, striatum and STN inputs are differentially integrated by the two subpopulations ^12^. In awake cats, intact striatal projections are necessary for the normal production of bursts in the GPe ^29^. Equally important, Aristieta *et al*. ^12^ demonstrated that the axon collateral inhibition from prototypic neurons powerfully shapes the activity of arkypallidal neurons, whereas the collateral connections from arkypallidal to prototypic neurons are sparse. Here, we did not observe any major deviation from independent activity in the CCHs/JPSTHs of LFD/HFD neurons (Figs. 5c,f and 6c, and Supplementary Figs. 6c,d and 7c). This lack of correlated activity between LFD/HFD neurons implies weak, if not nonexistent, functional connectivity between the LFD and HFD neurons during the processing of incentive cues. Further experiments in behaving animals should therefore be conducted to assess the functional connectivity between the two distinct pallidal subpopulations (NHP LFD/HFD neurons and rodent arkypallidal/prototypic neurons) during both motor and non-motor behaviors. Nevertheless, this study demonstrates a strong and sustained correlated activity in a NHP model between simultaneously recorded LFD neurons, and forces us to revise the view that all pallidal activity is independent (uncorrelated) in the healthy state ^42^.

### Are the NHP GPe LFD and HFD neurons the homologous rodent arkypallidal and prototypic neurons, respectively?

Arkypallidal neurons account for only one-fourth of all rodent GPe neurons and coexpress preproenkephalin (PPE) and the transcription factors FoxP2 and Meis2, whereas the prototypic neurons comprise approximately two-thirds of all GPe neurons and coexpress parvalbumin (PV) and transcription factors Nkx2-1 and Lhx6 ^10,11,14,43^. Remarkably, although we do not know whether LFD and HFD neurons exhibit distinct neurochemical features, the proportions of these neurons resemble those of the arkypallidal and prototypic neurons, respectively ^5,30^. In addition, NHP LFD/HFD neurons and arkypallidal/prototypic neurons showed similarities in their discharge rates and patterns across different brain states. The arkypallidal neurons had a lower and irregular firing rate that was significantly reduced during slow wave activity/natural sleep. By contrast, the prototypic neurons had a high and regular firing rate regardless of brain state (i.e., EEG slow wave activity/natural sleep or EEG spontaneous activation/awake state) ^10,11,14^. However, in anesthetized ^11^ and freely moving ^14^ dopamine-intact rats, the firing rate and regularity of the prototypic neurons during slow wave activity were significantly lower than those during cortical activation/awake states. A decrease in the activity of the HFD neurons during natural slow wave sleep or eye closure has also been reported in monkeys ^44,45^. The discharge of LFD neurons has not been studied; however, the sampling bias of extracellular recordings towards units with high discharge rates might exclude LFD neurons with zero/slow discharge rate during sleep. Although two distinct GPe populations with firing pattern characteristics similar to those reported in primates (Fig. 1b) ^5,30^ has been identified in freely moving rats ^4^, evidence for bursting and pausing activity in the arkypallidal and prototypic neurons, respectively, is still sparse ^14,46^.

Finally, the GPe LFD burst features might provide some possible hints as to the GPe homology question. It has been shown that the prototypic neurons project to the STN, BG downstream structures and to lesser extent to the striatum, whereas the arkypallidal neurons exclusively innervate the striatum and target both striatal projection neurons and interneurons ^10^. Earlier neuronal tract-tracing and in vitro electrophysiological studies using brain slices from adult guinea pigs have found that some GPe cells exhibiting low-threshold spike (LTS) burst features project to the striatum ^47,48^. Here, the NHP LFD neurons were found to fire accelerating-decelerating (parabolic) high frequency bursts of short durations (Supplementary Fig. 3a-c, top). This accelerating-decelerating burst pattern of the LFD neurons is similar to the LTS bursts, and in particular to those observed in the cells of the reticular/perigeniculate nucleus of the thalamus during slow wave sleep ^21,22^, which are known to be generated by hyperpolarization activation of transient low-threshold Ca^2+^ current (T current). The LTS burst features of the LFD neurons therefore constitutes another solid electrophysiological argument suggesting that the LFD neurons might be homologous to rodent arkypallidal neurons which exclusively project to the striatum.

Taken together, the quantitative analyses of the discharge features of both the NHP LFD and HFD neurons in the current study provide new findings that contribute to a better comparison of NHP LFD/HFD neurons and rodent arkypallidal/prototypic neurons. Nevertheless, the neuronal heterogeneity of the GPe might be more complex than the bipartite arkypallidal vs. prototypic organization ^15,43^. In particular, rodent GPe prototypic neurons have been genetically divided into Lhx6 and PV GPe neurons ^49^. The specific modulation of these two GPe neuronal subpopulations was shown to dramatically prolong the therapeutic benefits of deep brain stimulation in the 6-hydroxydopamine mouse model of Parkinson’s disease ^50^. Thus, further studies should aim to characterize the biochemical, structural and network features of NHP GPe LFD/HFD neurons, as well as examine the possible bursting/pausing capacity of rodent arkypallidal/prototypic neurons in details.

## Acknowledgments

We thank Dr. Yaron Dagan and Dr. Tamar Ravins for assistance with animal care, Dr. Atira Bick for assistance with the coordination and execution of the MRI, and Esther Singer for editing. We also thank Anatoly Shapochnikov for help in preparing the experimental set up and Dr. Hila Gabbay, Dr. Sharon Freeman and Dr. Uri Werner-Reiss for general assistance. This work was supported by the Nehemia Levtzion Fellowship and the Foulkes Foundation to S.K.; the French National Research Agency (ANR) and the French National Center for Scientific Research (CNRS) to M.D.; the Israel Science Foundation (ISF), the Israel-China BiNational Scientific Foundation and the TRR295 Rune German Science Centers Project, the Rosetrees and the Simone and Bernard Guttman Chair in Brain Research to H.B.

## Author contributions

S.K. and H.B., designed the research. S.K., A.A. and M.D. collected the data. S.K. analyzed and interpreted the data. S.K., M.D. and H.B. wrote the manuscript. All authors read and commented on the final version of the manuscript.

## Methods

### Animals and behavioral task

The experiments were conducted on four monkeys (Macaque fascicularis, 3 females R, 6 kg; Y, 3.5 kg; Le, 3 kg; and 1 male G, 4.5 kg) while they were engaged in an over-trained classical conditioning task with cues predicting reward (food), neutral or aversive (air-puff) outcomes (Fig. 1a). Different fractal cues (Chaos Pro 3.2 program, www.chaospro.de, displayed on a 17 inch LCD monitor, 50 cm in front of the monkey’s face) were presented to the monkey for a period of 2 s, and were immediately followed by an outcome signaled by one of three sounds that discriminated the possible events. All outcome deliveries were followed by a variable intertrial interval (5-6 s).

All experimental protocols were conducted in accordance with the National Research Council *Guide for the Care and Use of Laboratory Animals* and the Hebrew University for the use and care of laboratory animals in research and supervised by the Institutional Animal Care and Use Committee. The Hebrew University is an Association for Assessment and Accreditation of Laboratory Animal Care (AAALAC) internationally accredited institute.

### Recordings and data acquisition

After intensive training, a recording chamber was attached to the monkeys’ skulls. The recording chamber was tilted 40–50° laterally in the coronal plane and stereotaxically positioned to cover most of the GPe territory ^17,51^. The exact position of the chamber was established using a magnetic resonance imaging (MRI) scan and electrophysiological mapping.

Details of surgery, magnetic resonance imaging and data-recording methods are described in previous manuscripts ^30,36,52^. Briefly, during recording sessions, the monkeys’ heads were immobilized and eight glass-coated tungsten microelectrodes (impedance at 1000 Hz range: 0.2-0.8 MΩ) were advanced separately (EPS; AlphaOmega Engineering) toward and through the GPe. The electrical activity was amplified with a gain of 5K, band-pass filtered with a 1- 6000 Hz four-pole Butterworth filter and continuously sampled at 25 kHz by a 12-bit (± 5V input range) Analog/Digital (A/D) converter (MCP, AlphaOmega Engineering, Israel). Spike activity was sorted and classified online using a template-matching algorithm (ASD; AlphaOmega Engineering).

GPe neurons were identified according to the stereotaxic coordinates (based on the MRI and the primate atlas data) and real-time electrophysiological characteristics. Neurons exhibiting a low-frequency discharge with occasional brief high frequency bursts were identified online as LFD neurons ^5^, while neurons with a high-frequency discharge with recurrent pauses were identified as HFD neurons ^5,30^ (Fig. 1b). Recorded units were subjected to offline quality analysis that included tests for recording time duration, firing rate stability, refractory period and isolation quality. Firing rate was graphically displayed as a function of time and the longest stable recording segment was selected for further analysis while the rest of the data was discarded. Only neurons with a stable firing rate for ≥ 6 min and an isolation score ^53^ ≥ 0.6 were included in the database (n = 730 neurons). Table 1 provides the statistical details of the cells that were included in the analysis database.

**Table 1.** The neuronal database. n is the number of cells that met the inclusion criteria. The isolation score evaluates the quality of spike data and ranges from 0 to 1. Fraction ISI is the fraction of ISIs shorter than 2 ms out of all the ISIs of a cell. Recorded time stands for the recording period (in min) selected for analysis after visual assessment of the discharge rate stability for each neuron. Values are means ± SEM.

| Population | n | Isolation score | Fraction ISI<br>< 2 ms | Recorded time<br>(min) |
| --- | --- | --- | --- | --- |
| LFD | 98 | 0.82 ± 0.012 | 0.0203 ± 0.0035 | 35.11 ± 2.98 |
| HFD | 632 | 0.87 ± 0.004 | 0.0117 ± 0.0008 | 33.67 ± 1.05 |

### Discharge pattern assessment of spike train

For each spike train, we calculated the interspike intervals (ISI) and generated the ISI histograms of the well-isolated stationary units. In parallel, we computed the auto-correlograms of the spike train, calculated for ± 2500 ms offset with 1-ms bins.

We further examined the bursting and pausing features of the GPe neurons. For burst detection, we applied the Poisson surprise method with the surprise maximization (SM) search algorithm ^54^ to each spike train of the GPe neurons, using the following parameters: minimal burst length = 3 spikes; threshold surprise value (S) = 10; burst ISI limit = mean (ISI)/2; add limit = 150% of burst ISI limit and inclusion criteria (IC) = 5. Using these burst detection criteria, we calculated the number of spikes per burst and the burst durations, and characterized the intraburst ISI patterns (i.e., the ISI duration during the burst was further tested as a function of ISI ordinal in the burst).

For pause detection, we used the pause surprise method ^30^. This method is based on an evaluation of how improbable it is for a certain number of spikes or less to appear in a time segment of a spike train with a known average firing rate. Here, the algorithm began by detecting ISI longer than a certain length (minimal core interval). Then, it tested whether adding another neighboring ISI increased the surprise value. If so, the pause was redefined to include this ISI. Based on this procedure, pauses that did not exceed a certain length (minimal length of the final pause) were omitted. Additionally, pauses that were adjacent to one another were combined.

As in earlier studies ^30–32^, the parameters for the detection of the pauses in the HFD neurons were set as follows: minimal core interval = 250 ms, upper limit for added ISI = 5 (for each side of the core interval), minimal length of the final pause = 300 ms and maximum number of spikes enabling merging of pauses = 1. For the detection of pauses in the LFD neurons, some of the parameters were adjusted to account for the differences in the spike train between LFD and HFD neurons. Thus, since the mean ISI of the LFD neurons was 5.5 times longer than the mean ISI of the HFD (Supplementary Fig. 1b), the minimal core interval and minimal length of the final pause for pause detection in the LFD neurons were set to 1,375 (250 * 5.5) and 1,650 (300 * 5.5) ms, respectively.

Finally, we calculated the frequency and the mean duration of the bursts and pauses over their entire recording span. These two metrics were used to determine the burst/pause prevalence, which was defined as: frequency * mean burst/pause duration.

### Task-related neuronal responses

Neuronal responses to the different cue presentations (reward, neutral and aversive) were characterized by their peri-stimulus time histograms (PSTHs). The PSTHs (i.e., three PSTHs per neuron, 2 s each) were calculated in 1-ms bins and smoothed with a Gaussian window (SD = 20 ms). For each smoothed PSTH, we calculated the relative deviation from the baseline (i.e., relative PSTH) by subtracting the baseline firing rate (i.e., mean firing rate in the last 0.5 s of the intertrial interval) from the PSTH. The population response was estimated by averaging the relative PSTH deviations across the whole population. For each neuron, we also calculated the reward, neutral and aversive cue-related activities which were defined as the mean values of the baseline-corrected activity during the 2 s duration of reward, neutral and aversive cues, respectively. The response index was also defined for each cell as the absolute difference between reward or aversive cue-related activity and neutral cue-related activity ^23,24^. Signal correlation (SC) analysis ^25–27^ was then conducted to measure the similarity of the neural responses to the behavioral events of all pairs of (simultaneously and non-simultaneously) recorded neurons of each structure. The three smoothed PSTHs of a single neuron were combined into one matrix with rows for each behavioral event and columns for each 20-ms bin. Each column (time bin) was Z-normalized, and the matrix was flattened into a single vector representing the normalized PSTH vector of a neuron. For each pair of neurons, we computed their SC by calculating the correlation coefficient of the two normalized PSTH vectors. We applied Fisher’s Z-transform to the SC values ^55^ to calculate their mean, variance and skewness. The mean, variance and skewness of the population values were obtained by inverse Fisher Z-transform. The resulting SC values thus ranged from +1 (for highly correlated response profiles) through 0 (non-correlated response profiles) to -1 (anti-correlated response profiles).

### Cross-correlation analysis

Spike-to-spike synchronization (only between simultaneously recorded LFD/LFD, HFD/HFD, and LFD/HFD neuron pairs) was determined using cross-correlation analysis ^56^. Cross-correlation functions (or cross-correlograms, CCHs), often termed as noise correlations, measure the firing rate (or firing probability) of the target neuron around the time that the reference neuron fires and can thus reveal the functional connectivity between the simultaneously recorded neuron pairs (e.g., direct excitatory, inhibitory synapses or common synaptic inputs). Flat CCHs indicate that a functional connectivity between the neurons does not exist (independent activity), or that it is too weak to be detected by the finite duration of the recording of the spike trains ^56^. CCHs were calculated for ± 2500 ms offset with 1-ms bins over the trials (regardless the type of trial, i.e., reward, neutral and aversive trials) to provide the raw CCH. Finally, each raw CCH was corrected by subtracting its shuffle predictor (i.e., CCH after shuffling trials of the target neuron) to remove possible stimulus/cue-induced relationship and averaged over neuron pairs. For each category of neuron pair, we measured the firing rate of the mean corrected CCHs at time = 0 (T_0_ value). To assure that our results were not due to the small number of simultaneously recorded LFD/LFD neuron pairs (compared to the other categories of neuron pairs), we randomly selected (1000 times) equivalent numbers of HFD/HFD and LFD/HFD neuron pairs and re-measured the T_0_ value of the mean corrected CCHs.

For burst-to-burst, pause-to-pause and burst-to-pause synchronizations of LFD/LFD, HFD/HFD and LFD/HFD neuron pairs, respectively, the corrected CCHs were averaged over all LFD bursts or HFD pauses detected in the reference neuron and T_0_ value corresponded to the mean frequency of LFD bursts or HFD pauses at time = 0 (i.e., LFD burst or HFD pause initiation).

For burst-to-spike synchronizations of LFD/LFD and LFD/HFD and pause-to-spike synchronizations of HFD/HFD and HFD/LFD neuron pairs, the corrected CCHs were averaged over all LFD bursts or HFD pauses detected in the reference neuron and T_0_ value corresponded to the mean firing rate of LFD or HFD neurons at time = 0 (i.e., LFD burst or HFD pause initiation). Note that in these cases, for each LFD/LFD and HFD/HFD neuron pairs, bursts or pauses in the first unit (reference neuron) are used to measure the activity of the second unit (target neuron) and vice-versa, thus doubling the numbers of LFD/LFD and HFD/HFD neuron pairs.

### Joint PSTH analysis

To characterize the temporal dynamics of the spike-to-spike (noise) correlation during the behavioral cues, we calculated the joint PSTH (JPSTH) matrix ^25,26,28^. We first calculated the raw JPSTH, in which the (t_1_, t_2_) time bin was the count of the number of times there was a coincident event in which neuron number one spiked in time bin t_1_ and neuron number two spiked in time bin t_2_ in the same trial. To correct for rate modulations, we subtracted the shift predictor matrix (i.e., JPSTH after shifting all trials of neuron number two by one trial) from the raw JPSTH matrix. The JPSTH was calculated in bins of 50 ms and smoothed with a two-dimensional Gaussian window with an SD of 50 ms (single bin). Finally, we calculated the JPSTH main diagonals to quantify the time-depended modulation of zero lag (0-ms) noise correlation.

### Software and Statistics

All the data and statistical analyses were carried out using custom-made MATLAB 7.5 routines (Mathworks, Natick, MA, USA). Nonparametric statistical tests were used to determine significant differences between neuronal activities and cell properties. Mann-Whitney U-test/Kruskal-Wallis H-test and Wilcoxon signed rank/Friedman tests were used for statistical comparisons of unpaired and paired sample means, respectively. The criterion for statistical significance was set at p < 0.05 for all statistical tests. If necessary, a Bonferroni correction was used to adjust the chosen significance level according to the number of multiple comparisons.

---

**Supplementary Fig. 1.**
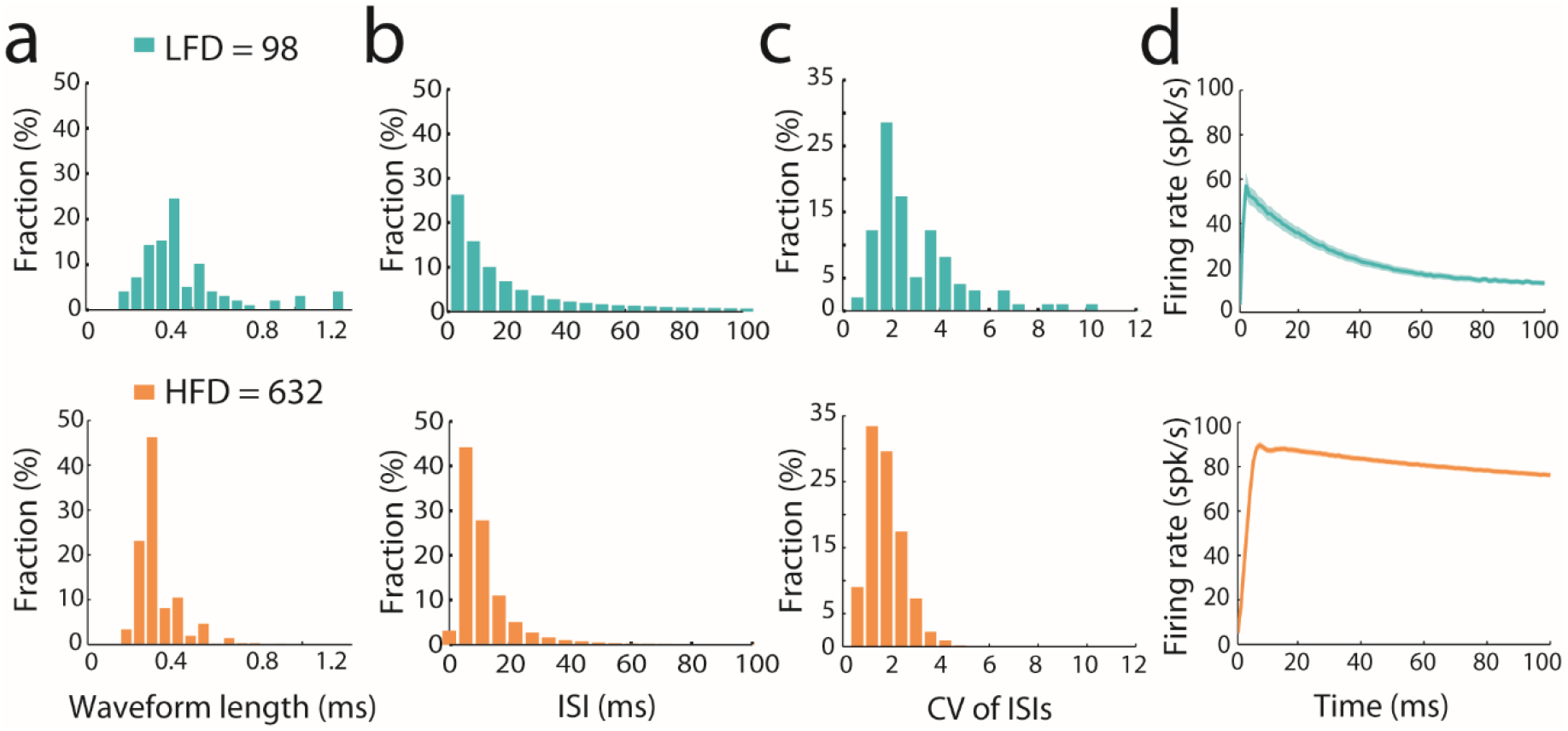
Electrophysiological properties of LFD vs. HFD neurons. **a**, Histograms of the waveform length distributions of LFD and HFD neurons (bin width = 0.1 ms). **b**, Interspike interval (ISI) histograms (bin width = 5 ms; n = 2,741,478 and 94,193,197 ISIs for LFD and HFD neurons, respectively). **c**, Histograms of the coefficient of variance (CV) of ISI distributions (bin width = 0.6). **d**, Mean auto-correlograms (bin width = 1 ms). The shaded areas indicate the SEMs.

**Supplementary Fig. 2.**
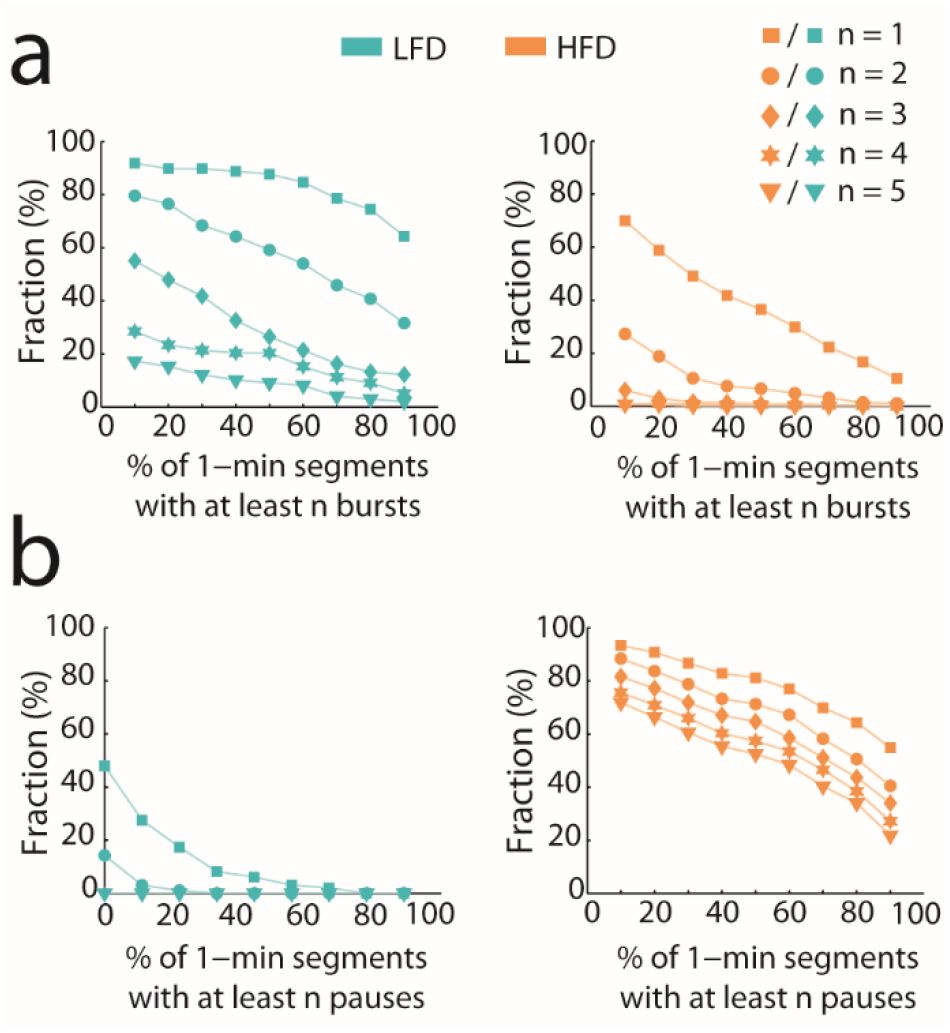
LFD tendency to burst vs. HFD tendency to pause. **a**, Fraction of LFD and HFD neurons with 10%-90% of 1-min segments of spiking activity with at least n = 1-5 bursts. **b**, Fraction of LFD and HFD neurons with 10%-90% of 1-min segments of spiking activity with at least n = 1-5 pauses.

**Supplementary Fig. 3.**
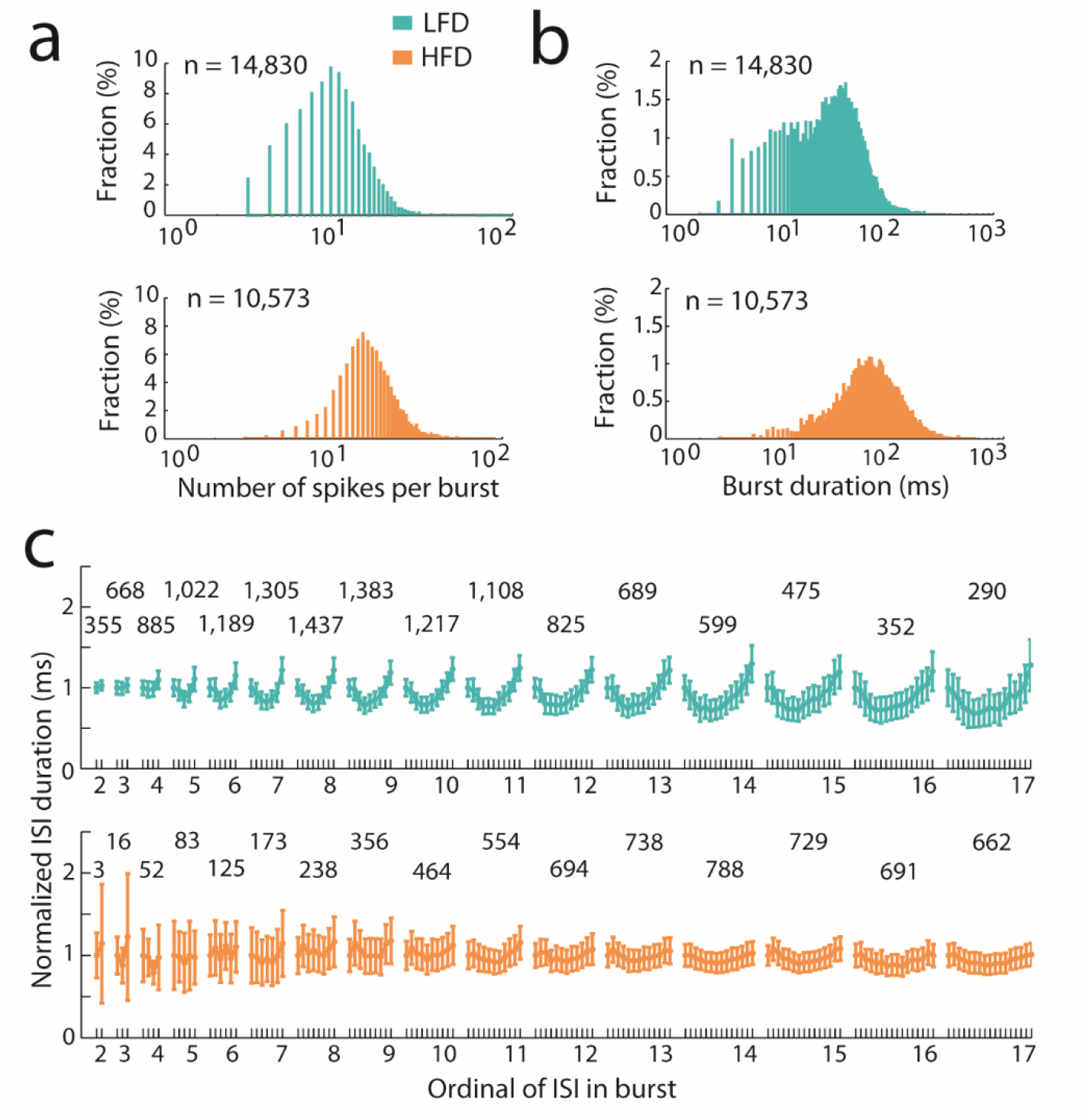
Burst properties of LFD vs. HFD neurons. **a**, Semi-log scale histograms of the number of spikes per burst (bin width = 1 ms) of LFD and HFD neurons. **b**, Semi-log scale burst duration histograms (bin width = 1 ms). **c**, Averaged intraburst interval pattern presented for bursts of different lengths. Results are presented as mean ± SEM values across all population bursts. Numbers above the plots indicate the number of observations for each burst length. LFD = 13,799 bursts; HFD = 6,366 bursts). LFD bursts ≥ 18 ISIs had fewer than 250 instances and are not presented (the same parabolic pattern was observed for these burst lengths). Same analysis was conducted on HFD neurons for comparison.

**Supplementary Fig. 4.**
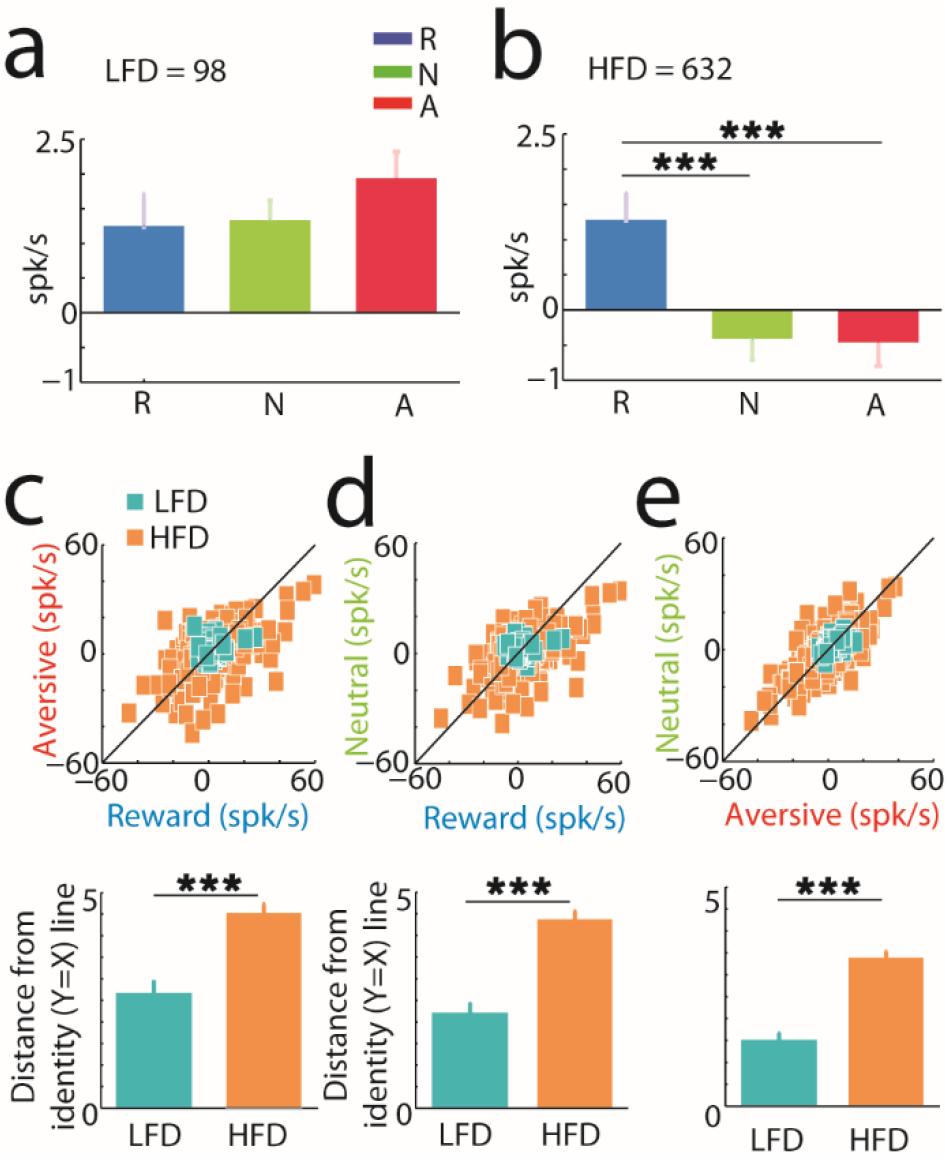
Early LFD and HFD neuronal responses encode the salience and valence of incentive cues, respectively. **a-b**, Mean LFD and HFD reward, neutral and aversive cue-related activities. Cue-related activity was defined as the mean value of the activity for a cue duration of 0.5 s (early neuronal response). R, responses to reward cues; N, responses to neutral cues; A, responses to aversive cues; Error bars represent SEMs; ^***^ indicates significant (p < 0.001) differences between cue-related activities. **c-e**, Top, comparison between reward/aversive, reward/neutral and aversive/neutral cue-related activities. Each square refers to a single neuron. The black line is the identity (Y = X) line. Bottom, mean distance values of the squares from the identity line. Error bars represent SEMs; ^***^ indicates significant (p < 0.001) differences between LFD and HFD neurons.

**Supplementary Fig. 5.**
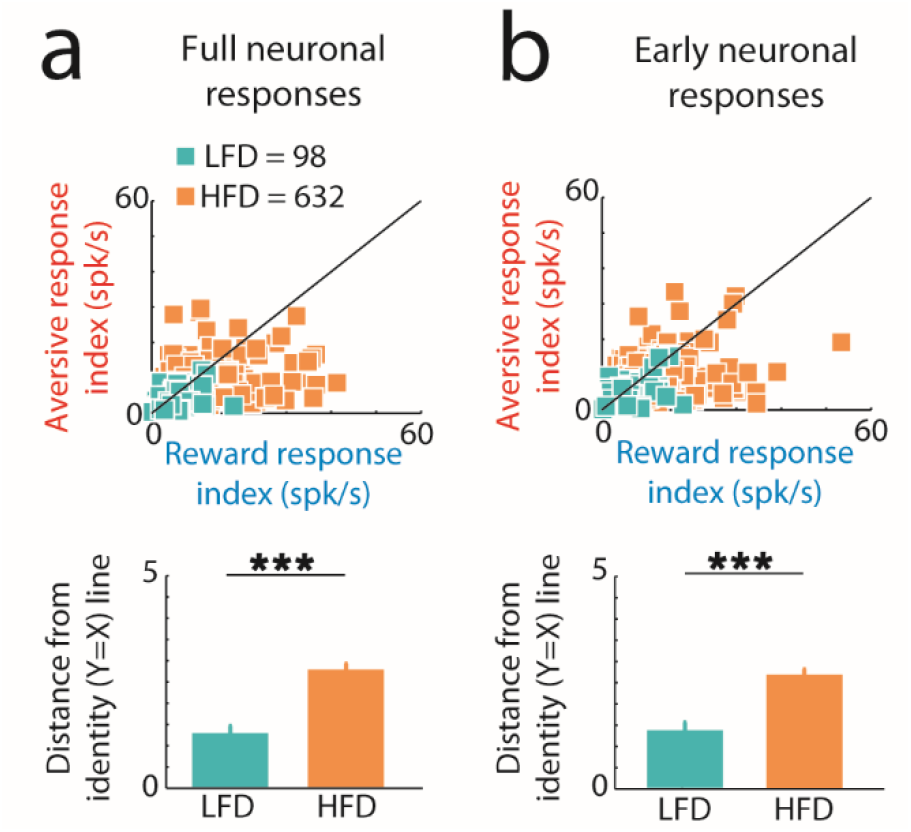
Encoding of reward and aversive cue values vs. the neutral cue value. **a**, Top, comparison of the response index of individual neurons to reward and aversive cues. The response index was calculated for each cell as the absolute difference between reward or aversive cue-related activity and neutral cue-related activity. Cue-related activity was defined as the mean value of the activity for a cue duration of 2 s (full neuronal response). Each square refers to a single neuron. The black line is the identity (Y = X) line. Squares below this line represent cells whose response index was larger for reward cues than for aversive cues. Bottom, mean distance values of the squares from the identity line. Error bars represent SEMs; ^***^ indicates significant (p < 0.001) differences between LFD and HFD neurons. **b**, Same conventions as in **a**, but cue-related activity was defined as the mean value of the activity for a cue duration of 0.5 s (early neuronal response).

**Supplementary Fig. 6.**
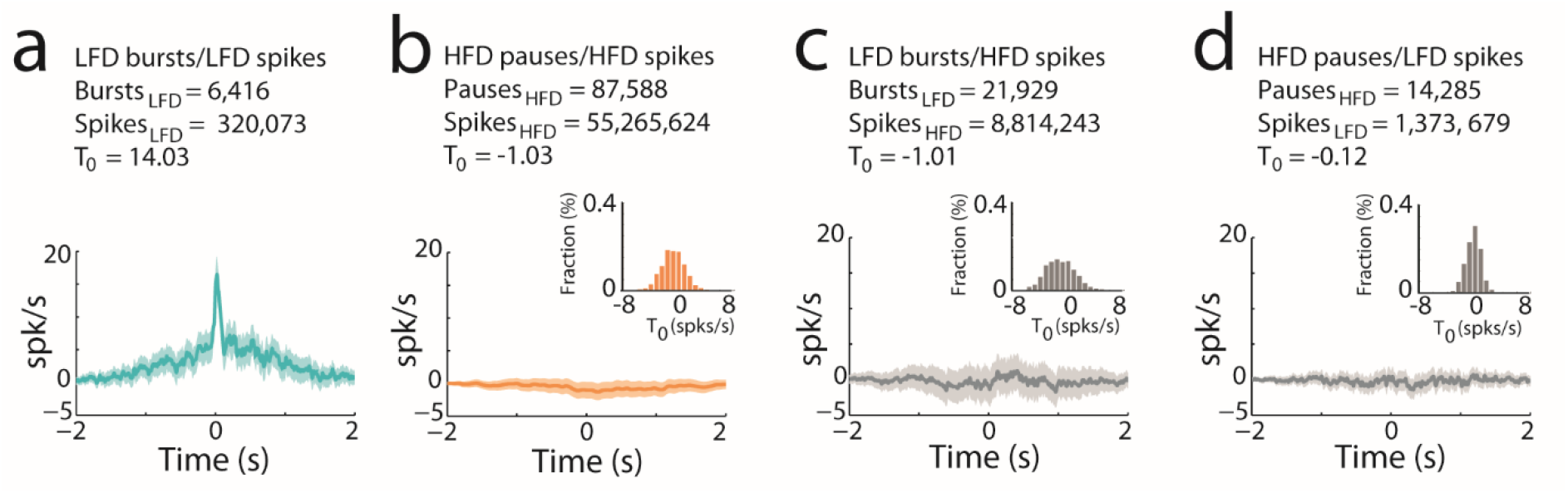
LFD bursts coincide with a sharp increase in the firing rate of simultaneously recorded LFD neurons. **a-d**, Average burst-to-spike, pause-to-spike, burst-to-spike and pause-to-spike cross-correlograms of LFD/LFD (n = 40), HFD/HFD (n = 1,398), LFD/HFD (n = 177) and HFD/LFD (n = 177) neuron pairs, respectively. Each cross-correlogram was corrected by subtracting its shuffled predictor in bins of 1 ms and then averaged over all LFD bursts or HFD pauses. For each LFD/LFD and HFD/HFD neuron pairs, bursts or pauses in the first unit were used to measure the activity of the second unit and vice-versa, thus doubling the numbers of LFD/LFD and HFD/HFD neuron pairs. The shaded areas mark SEMs; the T_0_ value is the mean firing rate of the LFD or HFD neurons at time = 0 (i.e., LFD burst or HFD pause initiation); insets are the distributions of the T_0_ values of the mean corrected cross-correlograms of 1000 randomly selected 40 HFD/HFD, LFD/HFD and HFD/LFD neuron pairs (to match the number of LFD/LFD neuron pairs).

**Supplementary Fig. 7.**
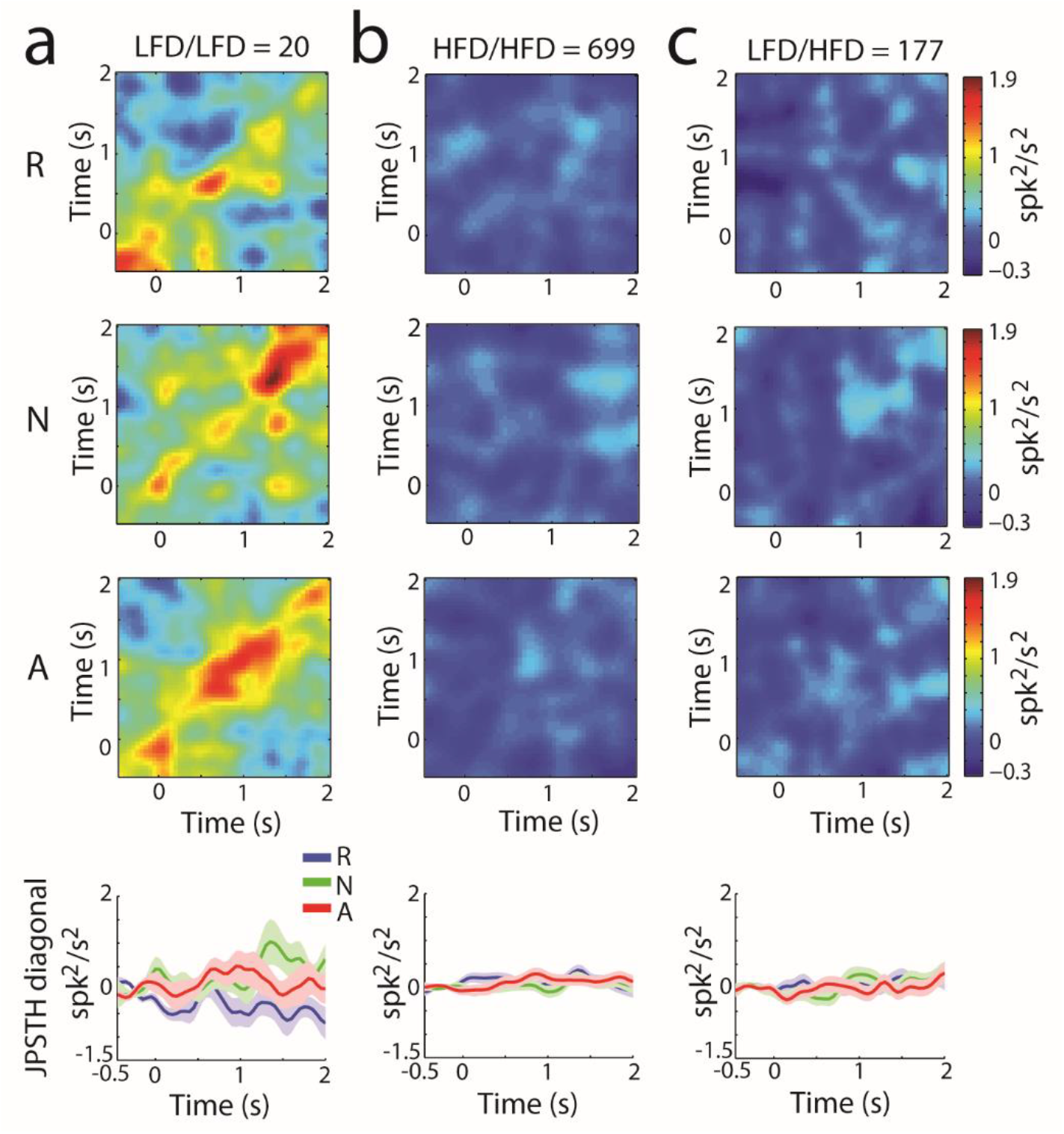
Dynamics of the noise correlations for LFD/LFD, HFD/HFD and LFD/HFD neuron pairs in different behavioral events. **a-c**, Population JPSTHs of LFD/LFD, HFD/HFD and LFD/HFD neuron pairs for reward (R), neutral (N) and aversive (A) cues presented at time 0. Each JPSTH was corrected by subtracting its JPSTH predictor in bins of 50 ms, smoothed with a two-dimensional Gaussian window (SD = 50 ms) and then averaged over all pairs. The different JPSTHs have the same color scaling (color bar on the right) to enable comparisons between behavioral trials. Bottom, JPSTH main diagonals (± SEM, shaded envelop) for LFD/LFD, HFD/HFD and LFD/HFD neuron pairs. The main JPSTH diagonal of each pair was normalized by subtracting the mean value of the last 0.5 s of the intertrial interval, and then averaged across the whole population. Blue, reward cues; Green, neutral cues; Red, aversive cues.

